# Prevalence and public health significance of Lyssavirus in bats in North region of Cameroon

**DOI:** 10.1101/2022.09.12.507528

**Authors:** Isaac Dah, Rodrigue Simonet Poueme Namegni, Moctar Mouiche Mouliom Mohamed, Simon Dickmu Jumbo, Ranyl Nguena Guefack Noumedem, Isabelle Conclois, Liegeois Florian, Laurent God-Yang, Jean Marc Feussom Kameni, Abel Wade, Dorothée Missé, Julius Awah-Ndukum

## Abstract

**Background:** Rabies is a zoonotic disease of all warm-blooded animals including humans. Though, there is little knowledge of the status of rabies in wild animals in Cameroon, the disease is endemic in the country with dogs being the main source of transmission. Bat habitats are widespread in Cameroon, but there is little information on the prevalence of rabies-like viruses in bats, nor the role of bats as a potential reservoir of rabies.

**Methods:** A cross sectional study was carried out to determine the prevalence and risk factors of Lyssavirus in bats in the Northern region of Cameroon. A total of 212 bats belonging to three families (Pteropodidae, Vespertilionidae, Molossidae) and 5 species were randomly sampled in 7 localities in the North of Cameroon and were tested for Lyssavirus antigen using direct Immunofluorescence Test (IFA). Overall, 57 (26.89%) of the bats collected showed an IFA positive reaction. The prevalence was higher (P<0.05) in adult bats (33.33%, 95% CI: 25.15 – 42.66) compared to young individuals (20.19%). The main risk factors identified in the study for human exposure to bats were gender (Male), educational level (tertiary), religion (Christianity), ethnic group (Matal), the presence of bats in the area, the practice of bat hunting and consumption and the level of awareness on bat rabies-like viruses.

**Conclusion:** The study found the first evidence of Lyssavirus in bats in Cameroon. This finding revealed that bat rabies-like viruses are real and constitutes a potential human health problem in communities with bat habitats in the North region of Cameroon. Enhancing the level of public awareness and health education on the potential of bats as reservoirs of Lyssavirus in Cameroon as well as the integration of the “One Health” approach for effective management of animal and human rabies should be emphasized.

**Author summary:** Rabies is a zoonotic disease caused by a virus of the genus Lyssavirus. It affects all warm-blooded animals including humans. Canine and Human rabies are well documented as endemic in Cameroon, but little is known about this disease in wildlife, in particular among bats, despite their multiple interactions with the inhabitants of Northern Cameroon. Indeed, bats were hunted, sold and eaten as bush meat by local populations. We investigated the presence of Lyssavirus in bat and assessed the risk factors of human exposure to bats in the Northern region of Cameroon. The study highlights that Lyssavirus is present in bats in this area. The population was aware of human and canine rabies, however, the presence of the disease in bats was less known. Based on these findings, investigating bat populations on a large scale, to characterise the Lyssavirus strains circulating in the region, as well as educate the local population on the risks of rabies transmission from bats to humans and other animals.

## Introduction

Rabies is a virulent fatal zoonotic disease of major public health concern caused by a Lyssavirus [1]. The virus is known to affect all warm-blooded animals, including domestic pets, which are the main vectors. Bat Lyssaviruses are distributed worldwide including canine rabies-free countries and bats are the primary dispensers of Lyssavirus to other animals and humans over vast geographical areas through their saliva and urine [2]. Bats have been directly involved in the transmission of rabies to humans through aerosols and unapparent bites as well as indirectly through animals they infected Infections of others animal could be through contact with contaminated body fluids by scratching of mucous membranes and open skin wounds [3, 4, 5, 6, 7]. Fatalities in humans and other animal species due to bat Lyssavirus [8, 9, 10], as well as human rabies related to Lyssavirus bat variants, have been reported to be associated with bat biting and simple contact with bats [11]. Bat Lyssaviruses are more infectious in superficial epidermal inoculation and multiply faster in non-nerve cells at a lower temperature than the canine rabies virus [12]. The transmission of animal rabies to humans is higher in areas where animal rabies is widespread [13], and there are frequent human and animal exposures to bats, including sick and injured bats with high risk of contamination [14].

Rabies has induced over 3.7 million (95% CI: 1.6 – 10.4 million) disability-adjusted life years (DALYs), 8.6 billion USD (95% CI: 2.9 – 21.5 billion) economic losses annually and 59,000 human deaths per year worldwide [15]. Bat Lyssavirus could be underestimated in Africa, due to insufficient surveillance programs [16]. The disease has been reported in bat populations in parts of the continent with prevalence ranging from 29 to 67% in Kenya [17], 38% in Ghana [18], 19% in Nigeria [19]) and 5.5% in the Democratic Republic of Congo [20]. In Cameroon, over US $ 576,232.88 was estimated as direct financial losses linked to rabies prevention measures and post-exposure treatments between 2004 and 2013 in three cities (Garoua, Yaoundé and Ngaoundéré) of Cameroon [21]. Urban rabies is widespread in dogs, which are considered as the main source of animal and human rabies in Cameroon [21, 22, 23, 24, 25] and canine and human rabies are endemic in the Northern region of the country [21, 23, 26]. There are many bat colonies and a majority of bat roosting sites are located adjacent to human communities where the level of physical interactions between bats and humans is high in many parts of the country. However, there is little or no attention on the epidemiology of bat Lyssaviruses. Insectivorous bats also frequently flock and inhabit roof-tops and homes while hunting of frugivorous bats for food are common in Cameroon. According to Mickleburgh *et al*. [27], the consumption of these animals by local populations could endanger the long-term survival of these species (*Eidolon helvum*). Indeed, *E, helvum* is consumed and marketed locally in the Bomboko area (South-West) and elsewhere in Cameroon, where they constitute an important source of income for local hunters [27]. Meanwhile, human–wildlife interactions that increase the risk of transmission are frequent and various, namely hunting, butchering and consuming wild animals, including bats, are common in Cameroon [28]. Additionally, subsistence activities and large-scale agriculture expose people not only to bat bites, but to potential infection through scratching due to the presence of urine and droppings of bats [28]. In Northern Cameroon, bats usually form colonies in the roofs, trees and abandoned tall structures in human communities and are also hunted as a source of animal protein and guano, which includes bat droppings, are widely collected as fertilizer [29]. In this context, the present study was carried out to determine the prevalence of rabies related-virus in bats and associated risk factors to human health in the Northern region of Cameroon.

## Material and Methods

### Description of study areas

This study was carried out in 7 localities (Babla, Djalingo, Lagdo, Guider, Mayo Oulo, Yelwa and Garoua II) in two Administrative Divisions (Mayo-Louti and Bénoué) of the North region of Cameroon (6° - 10° N and 12° - 16°E) (Fig 1) based on the available information on bat roosts, bat colonies and field observations of flying and foraging bats. The North region is situated in the Sudano-sahelian region with an average altitude of 249 m, a short rainy season from mid-March to October of 1200 – 1600 mm per annum and an ambient temperature range from 21° to 36°C.

**Fig 1:**
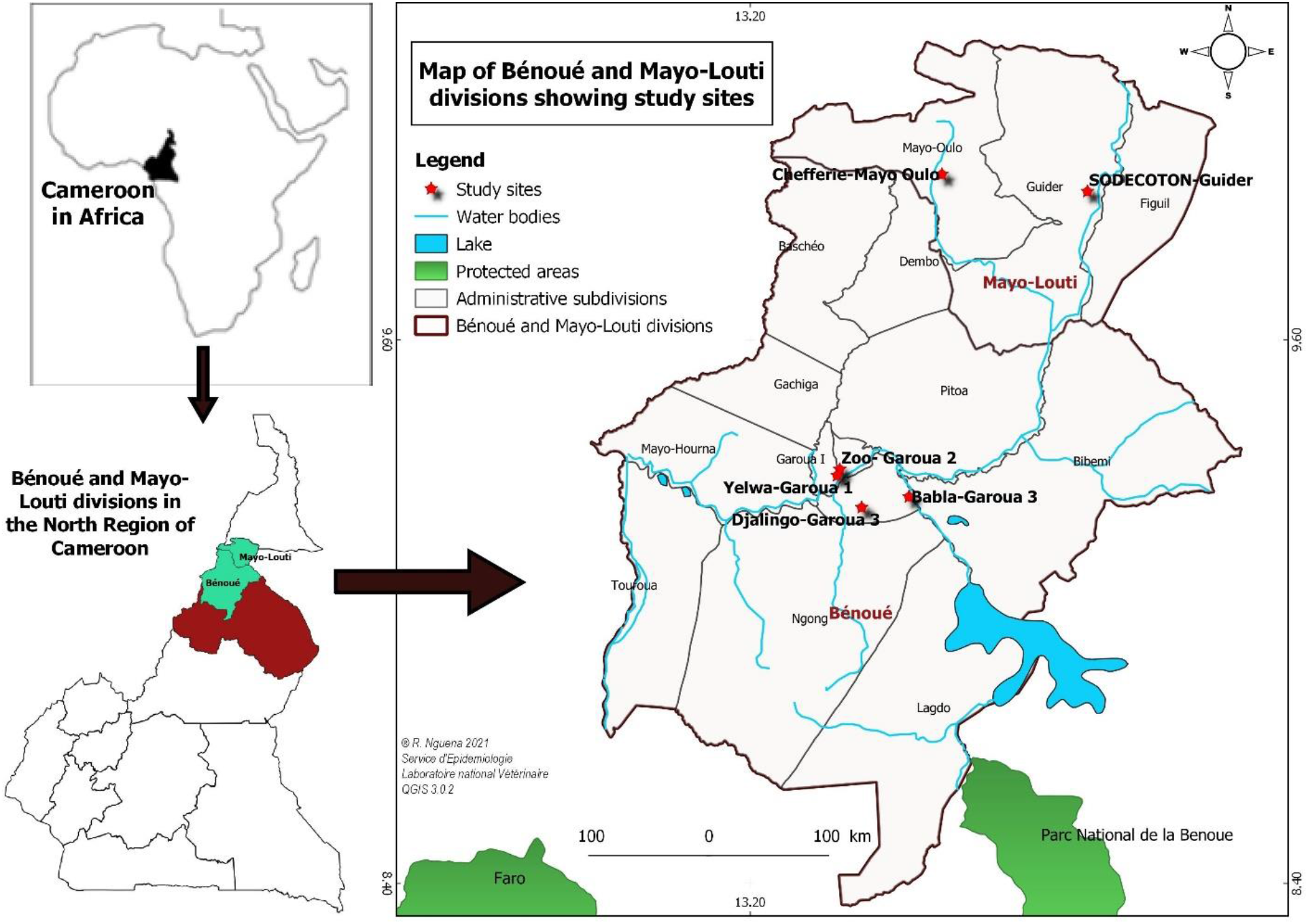
Map showing study sites. Bats specimen were collected in several study sites (red stars) in Mayo-Louti and Bénoué administrative divisions of the North Cameroon region.

### Sampling of bats for the study

A cross sectional study was conducted in the North Region of Cameroon from February to May 2017 following identification and listing of communities and geographical areas with bats roosts and bat activities. Information on bats roosts was obtained through the aid of local communities’ leaders. All identified localities with bat activities (hunting, selling of bat as bush meat etc.) were included and visited for sample collection in the study. Insectivorous bats and Fruit bats were collected twice weekly from the hunters in exchange of money before delivery to their clients. Indeed, there are vendors who bought bats and resold the meat to the consumers in these localities. Also, 7 dead bats including 6 *Eidolon helvum* at the Garoua zoological garden and 1 *Chaerephon pumilus* at Lagdo were also collected during the study period. Whole animals were shipped to the National Veterinary Laboratory (LANAVET) Garoua in an individual ziplock bag placed in a cooler with frozen ice pack [30]. The sample size was determined according to Thrusfield [31] based on a bat Lyssavirus prevalence of 19% obtained in Nigeria [19]. Overall, 212 collected from hunters and dead bats made up of 76 frugivorous bats (*Eidolon helvum*) and 136 insectivorous bats (10 *Chaerephon chapini;* 10 *Chaerephon leucogaster;* 96 *Chaerephon pumilus* and 20 *Scotophilus leucogaster*) were collected in 7 localities of the North region.

### Laboratory analysis

Laboratory analysis was done at LANAVET Garoua, in the North Region of Cameroon. The bat species was identified based on biomorphometric measurements with calliper using dichotomous keys [29]. The sex of bats was determined based on the observation of external genital organs and stage of growth (whether the bats were juvenile or adult) was through appraisal of body development and pelage coloration as previously described [19, 32].

A cross-section of the brain (including cerebral cortex and cerebellum) of each bats collected (212) in the study was taken after skull dissection in a certified biosafety cabinet [33]. An equally portion of each brain part collected was mixed and a thin smear spread on a slide was made. The slides were fixed in cold acetone at −20°C for one hour and then dried for 30 min at room temperature. The Lyssavirus ribo-nucleoprotein complex was detected using Rabies specific labelled polyclonal antibodies with Evans blue (1/2000 final dilution) as counterstain to denote stained areas and incubated at 37°C in a humidified chamber according to the manufacturer’s instructions (Bio-Rad kit 357-2112, Marnes-La-Coquette, France). The direct Immunofluorescence Assay (IFA) was used to detect Lyssavirus antigen in brain tissue, in accordance with WOHA (World Organization for Animal Health) and WHO (World Health Organisation) recommendations [34]. Briefly, positive and negative canine rabies controls available at the National Veterinary Laboratory in Garoua were used to validate the test results. A test was validated when there was fluorescence in the positive control and no fluorescence in the negative control. Three experienced staff members read the test slides before validating the results.

### Analysis of bat rabies exposure risk factors

For risk factors to determine human exposure, a structured questionnaire was issued to 535 willing inhabitants around the bat collection sites. Briefly, households within 5km radius of the bats collection sites and present during a visit were randomly surveyed by simple number generation without replacement. An oral consent of each respondent was obtained in advance to participate in the study. Participants (≥15 years old) were interviewed individually to avoid communication bias during the survey. The questionnaires were structured to collect information on a range of variables including lifestyle, socio-demographic data, knowledge about rabies, bat activities and human – bat interactions.

### Statistics

Data obtained in the study was entered into Microsoft Excel (Microsoft, PC/windows XP, 2010, Redmond WA, USA) for descriptive statistics and transferred to IBM^®^ Statistical Package for Social Sciences Software (SPSS Inc., Chicago IL, USA) version 21 for further analysis. Variables of interest were compared between exposed and non-exposed persons using odds ratios (OR) with 95% confidence intervals (CI) and p-values were calculated with the chi square test or Fisher’s exact test where appropriate [35]. Exposure to bat Lyssavirus was defined as having been bitten by a bat, scratched by a bat or touched a bat with bare hands [36, 35] as well as having hunted bats, prepared and / or consumed bat meat. The presence of bat in living space (home, roofs and trees in vicinity of home) and / or contact with bat wastes (body fluid, faeces, urine) was also considered as risk exposure to bats.

## Results

### Prevalence of rabies in bats

Between February to May 2017, brain tissues samples were taken from 212 bats belonging to 3 families (Vespertilionidae, Molossidae, Pteropodidae), including 5 species (*Chaerephon chapini, Chaerephon leucogaster, Chaerephon pumilus, Eidolon helvum, Scotophilus leucogaster*). The tissue samples were tested using direct Immunofluorescence Assay (IFA) to detect the Lyssavirus ribo-nucleoprotein complex (Fig. 2 and Table 1).

**Table 1.** Prevalence of Lyssavirus in bats by Immunofluorescence Assay in North Cameroon according to endogenous and exogenous factors.

| Parameter | Variable | N | N+ | Prevalence (95% CI) | P-value (X<sup>2</sup> value) |
| --- | --- | --- | --- | --- | --- |
| **Families** | *Vespertilionidae* | 20 | 6 | 30 (14.55 – 51.90) | 0.9130 (0.1820) |
|  | *Molossidae* | 116 | 30 | 25.86 (18.76 – 34.51) |  |
|  | *Pteropodidae* | 76 | 21 | 27.63 (18.84 – 38.57) |  |
| **Species** | *Chaerephon chapini* | 10 | 2 | 20 (5.67 – 50.98) | 0.9625 (0.604) |
|  | *Chaerephon leucogaster* | 10 | 2 | 20 (5.67 – 50.98) |  |
|  | *Chaerephon pumilus* | 96 | 26 | 27.08 (19.20 – 36.72) |  |
|  | *Eidolon helvum* | 76 | 21 | 27.63 (18.84 – 38.57) |  |
|  | *Scotophilus leucogaster* | 20 | 6 | 30 (14.55 – 51.90) |  |
| **Sex** | Male | 88 | 26 | 29.54 (21.03 – 39.78) | 0.4620 (0.54099) |
|  | Female | 124 | 31 | 25.00 (18.21 – 33.29) |  |
| **Age** | Adult | 108 | 36 | 33.33 (25.15 – 42.66) | 0.03098* (4.654) |
|  | Juvenile | 104 | 21 | 20.19 (13.60 – 28.90) |  |
| **Body condition score** | 1 (good) | 104 | 32 | 30.77 (22.72 – 40.19) | 0.3002 (2.4065) |
|  | 2 (medium) | 105 | 25 | 23.81 (16.68 – 32.79) |  |
|  | 3 (thin) | 3 | 0 | 0 |  |
| **Trophic diet** | Frugivorous | 76 | 21 | 27.63 (18.84 – 38.57) | 0.8549 (0.0334) |
|  | Insectivores | 136 | 36 | 26.47 (19.78 – 34.46) |  |
| **Samples collection site** | Babla (church) | 20 | 5 | 25 (11.19 – 46.87) | 0.6020 (4.5550) |
|  | Djalingo (church) | 45 | 9 | 20 (10.90 – 33.82) |  |
|  | Guider/Mayo Oulo | 6 | 3 | 50 (18.76 – 81.24) |  |
|  | Lagdo (Integrated Health centre of Plateau) | 55 | 17 | 30.82 (18.68 – 43.11) |  |
|  | Yelwa (primary school) | 16 | 3 | 18.75 (6.59 – 43.01) |  |
|  | Zoological garden of Garoua | 70 | 20 | 28.57 (17.98 – 39.15) |  |
|  | Global prevalence |  |  | 26.89 (21.37 – 33.23) |  |

\* : P < 0,05 ; N: Number of samples tested, N+: Number of positive samples; () : Chi-2 value, IFA : Immunofluorescence Assay;

Out of 212 (26.89% (95% CI: 21.37 – 33.23) bat brain samples, 57 tested positive for Lyssavirus antigen using direct Immunofluorescence Assay (IFA) (Table 1).

**Fig 2.**
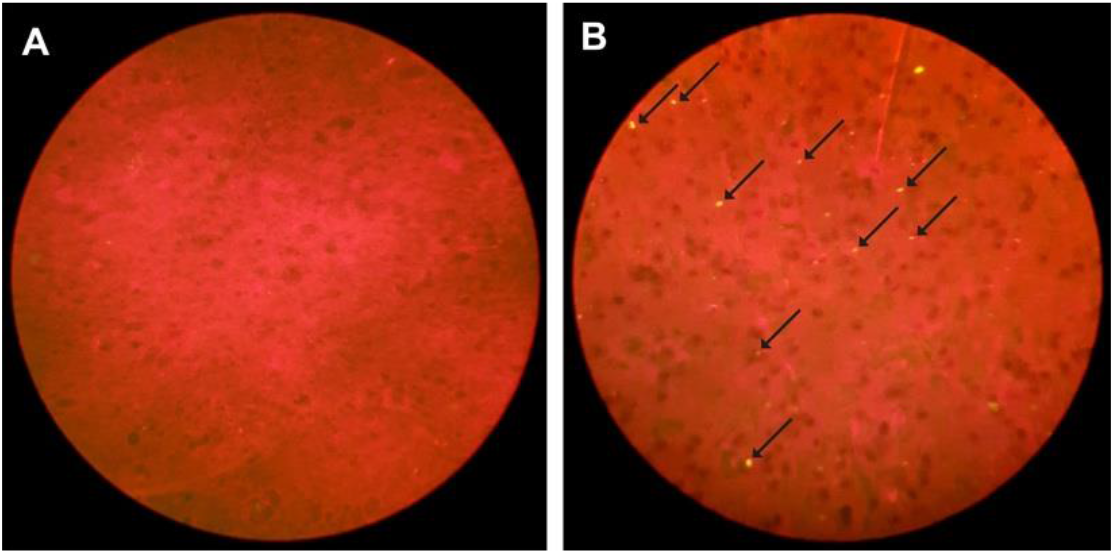
Detection of Rabies’ ribo-nucleoprotein complex using Direct Immunofluorescence Assay. Negative **(A)** and Positive **(B)** brain samples were fixed, analysed for the presence of the Rabies virus antigen and visualized under fluorescence microscope (100X objective). Arrows indicate viral ribo-nucleoprotein complex. Evans blue was added to the conjugate as a counterstain, which turned the tissue noticeably red to denote the green fluorescence.

The prevalence of Lyssavirus in bats in North Cameroon according to endogenous and exogenous factors is detailed in Table 1. The positivity rate was significantly higher in adult compared to young bats (p-value= 0.03002). The Lyssavirus prevalence in this study was not associated with bat families (p-value= 0.9130), bat species (p-value = 0.9625), sex (p-value =0.4620), body condition score (p-value =3002), trophic diet (p-value =0.8549) and collection sites (p-value =0.6020). There was at least one positive case of Lyssavirus in each of the bat families, each bat species tested and each collection site suggesting that the virus is circulating in bats in the region.

### Risk factors to human health of rabies

There were 535 volunteers of different religions, genders, and educational backgrounds that were interviewed. Data on a range of variables, including lifestyle, socio-demographics, and knowledge of rabies, bat activities, and human-bat interactions were collected.

The study showed more respondents were aware of canine rabies (74.57% (95%CI: 70.72 – 78.09)) than of bat rabies related viruses (4.67% (95%CI: 3.18 – 6.80)) and their zoonotic risks (Table 2). The proportion of respondents with knowledge of rabies was significantly influenced (P<0.05) by gender and level of education for canine rabies compared to level of education for bat rabies related virus. Overall, more male than female respondents showed higher (P<0.05) levels of awareness of canine rabies. Additionally, more literate respondents showed higher (P<0.05) levels of awareness of bat rabies than illiterate respondents (Table 2).

**Table 2.** Level of canine and bat rabies awareness of respondents according to socio-demographic characteristics in the Northern Region of Cameroon (n=535).

| Parameter / variable | N | Canine Rabies |  |  | Bat Lyassavirus |  |  |
| --- | --- | --- | --- | --- | --- | --- | --- |
| | | Yes (n) | Proportion (% [CI 95%]) | P-value ($X^2$) | Yes (n) | Proportion (% [CI 95] %) | P-value ($X^2$) |
| **Localities** |  |  |  |  |  |  |  |
| Babla | 87 | 63 | 72.41 (62.22 – 72.41) | 0.85 (2.63) | 6 | 6.90 (3.20 – 14.24) | 0.69 (3.92) |
| Djalingo | 62 | 46 | 74.19 (62.12 – 83.44) |  | 4 | 6.45 (2.54 – 15.45) |  |
| Garoua II | 70 | 53 | 75.71 (64.50 – 84.24) |  | 2 | 2.86 (0.79 – 9.84) |  |
| Guider | 107 | 77 | 71.96 (62.80 – 79.60) |  | 6 | 5.61 (2.60 – 11.70) |  |
| Lagdo | 70 | 55 | 78.57 (67.61 – 86.56) |  | 1 | 1.43 (0.25 – 7.66) |  |
| Mayo Oulo | 78 | 56 | 71.79 (60.96 – 80.57) |  | 3 | 3.85 (1.32 – 10.71) |  |
| Yelwa | 61 | 49 | 80.33 (68.69 – 88.37) |  | 3 | 4.92 (1.69 – 13.49) |  |
| **Age (years)** |  |  |  |  |  |  |  |
| [15 – 25] | 203 | 138 | 67.98 (61.28 – 74.01) | 0.07 (8.70) | 7 | 3.45 (1.68 – 6.95) | 0.42 (3.88) |
| [26 – 35] | 134 | 102 | 76.12 (68.24 – 82.55) |  | 8 | 5.97 (3.06 – 11.34) |  |
| [36 – 45] | 92 | 74 | 80.43 (71.17 – 87.25) |  | 5 | 5.43 (2.34 – 12.09) |  |
| [46 – 55] | 48 | 37 | 77.08 (63.46 – 86.69) |  | 4 | 8.33 (3.29 – 19.55) |  |
| [56 – 65] | 58 | 48 | 82.76 (71.09 – 90.36) |  | 1 | 1.72 (0.30 – 9.13) |  |
| **Religion** |  |  |  |  |  |  |  |
| Christians | 348 | 264 | 75.86 (71.10 – 80.06) | 0.59 (1.05) | 21 | 6.03 (3.98 – 9.04) | 0.05 (5.89) |
| Muslims | 158 | 115 | 72.78 (65.36 – 79.12) |  | 2 | 1.27 (0.35 – 4.50) |  |
| Without religion | 29 | 20 | 68.97 (50.77 – 82.73) |  | 2 | 6.90 (1.91 – 21.97) |  |
| **Gender** |  |  |  |  |  |  |  |
| Female | 209 | 146 | 69.86 (63.33 – 75.68) | 0.045* (4.04) | 6 | 2.87 (1.32 – 6.12) | 0.11 (2.50) |
| Male | 326 | 253 | 77.61 (72.78 – 81.80) |  | 19 | 5.83 (3.76 – 8.92) |  |
| **Educational level** |  |  |  |  |  |  |  |
| None | 136 | 91 | 66.91 (58.63 – 74.26) | <0.01* (12.74) | 2 | 1.47 (0.40 – 5.20) | <0.001* (17.65) |
| Primary | 151 | 111 | 73.51 (65.95 – 79.90) |  | 3 | 1.99 (0.60 – 5.68) |  |
| Secondary | 218 | 168 | 77.06 (71.04 – 82.14) |  | 15 | 6.88 (4.21 – 11.04) |  |
| Tertiary | 30 | 29 | 96.67 (83.33 – 99.41) |  | 5 | 16.67 (7.34 – 33.57) |  |
| **Total** | **535** | **399** | **74.57 (70.72 – 78.09)** |  | **25** | **4.67 (3.18 – 6.80)** |  |

N: number of participants, Yes: number of respondents who are awareness of canine and bat rabies; \*: $p<0.05$ ; (): value of Khi-2.

Respondents who were aware of canine rabies cited dogs (73.64%) and cats (19.81%) as the principal sources of rabies infection in humans (63.73%) and other animals. The cited modes of transmission were through bites (92.14%) and scratches (4%) of rabid animals as well as food (4%) and water (1.20%) contaminated by fluid and wastes from rabid animals. Goats (10.65%), cattle (7.85%), horses (7.1%), sheep (6.91%), pigs (5.42), fowl (4.48%), donkeys (0.25%), monkeys (0.25%), gorillas (0.25%) and mice (0.002) were also listed as susceptible animals and sources of rabies for humans and other animals (dogs, cats, cattle, sheep, goat, horse, donkey etc.)

Gender, educational level, religion and ethnicity of respondents significantly influenced (P<0.05) the level of exposure to bats and bat products in the Northern Region of Cameroon (Table 3). Similar to the level of rabies awareness among the respondents, more male and literate respondents and Christian respondents showed more (P<0.05) interaction with bats and bat products than female, illiterate and non-Christian respondents. The highest level of exposure was reported among the Matal community (83.33%; 95% CI: 55-19 – 95.30) while the lowest was reported among the Peulh community (18.18%; 95% CI: 7.31 – 38.51)) (Table 3).

**Table 3:**
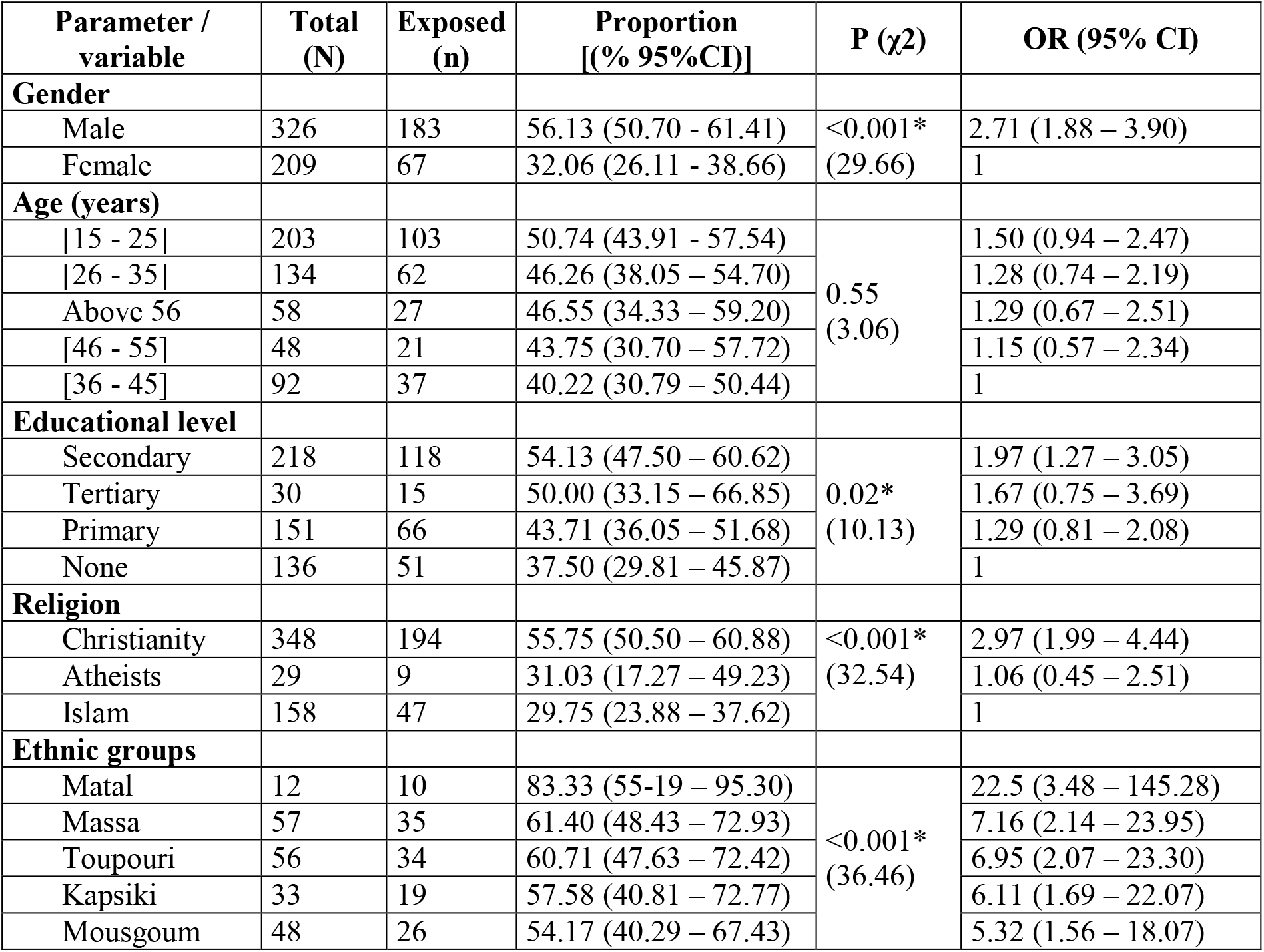
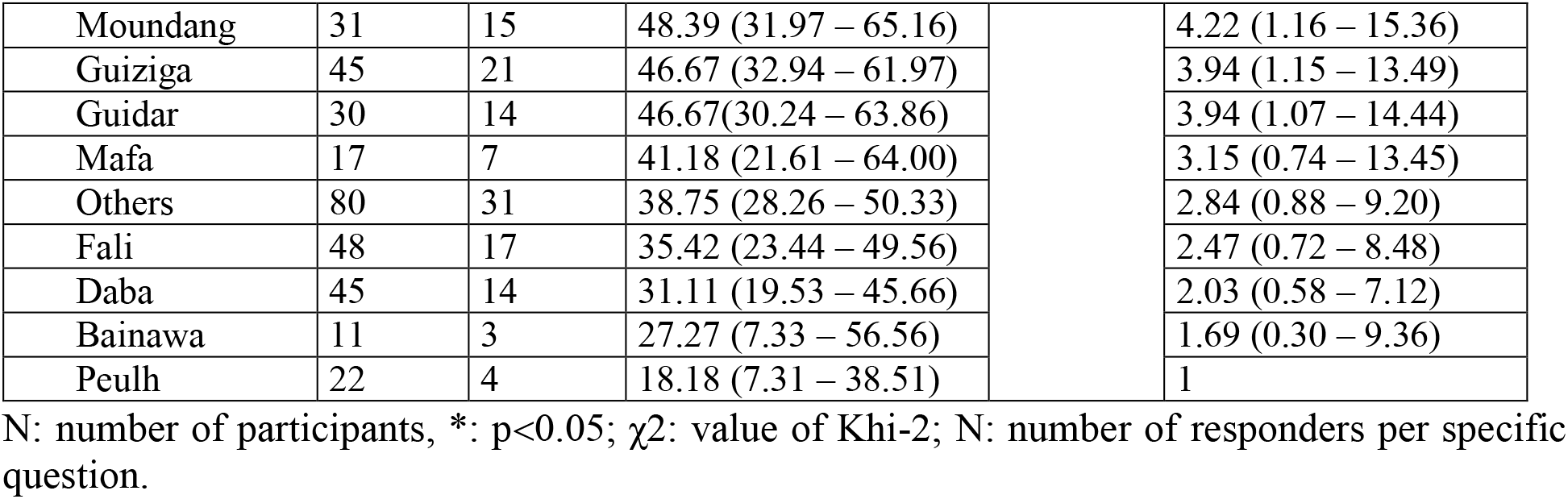
Risk factors associated with human exposure to bats and bat products in the North Region of Cameroon (n=535)

Though the respondents who were aware of canine rabies were significantly (p<0.05) more exposed to bat Lyssavirus than those who were less aware of the disease, there was no difference (P>0.05) in terms of exposure to bat Lyssavirus between respondents who were aware of zoonotic rabies and, in particular, bat rabies (Table 4). The presence of bats within a locality and homes, bat consumption, bat hunting and other interactions with bats significantly (P<0.05) exposed respondents to zoonotic rabies. The proportion of exposed respondents varied significantly (p<0.05) according to locality.

**Table 4:** Risks associated to exposure to bat rabies among respondents in the Northern Region of Cameroon.

| Parameter / variable | Total<br>N | Exposed<br>(n) | Proportion<br>% (95%CI) | P-value<br>($\chi^2$) | OR (95%CI) |
| --- | --- | --- | --- | --- | --- |
| **Awareness of canine rabies** |  |  |  |  |  |
| Yes | 399 | 198 | 49.62 (44.74 – 54.51) | 0.02*<br>(5.28) | 1.59 (1.07 – 2.37) |
| No | 136 | 52 | 38.24 (30.50 – 46.62) |  | 1 |
| **Aware about risk of zoonotic rabies** |  |  |  |  |  |
| No | 56 | 28 | 50.00 (37.33 – 62.67) | 0.95<br>(0.004) | 1.02 (0.58 – 1.79) |
| Yes | 341 | 169 | 49.56 (44.29 – 54.99) |  | 1 |
| **Aware about risk of bat rabies** |  |  |  |  |  |
| Yes | 25 | 16 | 64.00 (44.52 – 79.75) | 0.16<br>(1.98) | 1.82 (0.78 – 4.21) |
| No | 372 | 184 | 49.46 (44.41 – 54.52) |  | 1 |
| **Presence of bats in the locality** |  |  |  |  |  |
| Yes | 490 | 240 | 48.98 (44.58 – 53.40) | <0.001*<br>(11.85) | 3.36 (1.63 – 6.94) |
| No | 45 | 10 | 22.22 (12.54 – 36.27) |  | 1 |
| **Presence of bats in home** |  |  |  |  |  |
| Yes | 227 | 129 | 56.83 (50.33 – 63.11) | <0.001*<br>(16.16) | 2.03 (1.43 – 2.88) |
| No | 308 | 121 | 39.29 (34.00 – 44.84) |  | 1 |
| **Bat hunting** |  |  |  |  |  |
| Yes | 150 | 107 | 71.33 (63.63 – 77.96) | <0.001*<br>(50.69) | 4.21 (2.79 – 6.34) |
| No | 385 | 143 | 37.14 (32.46 – 42.20) |  | 1 |
| **Bat consumption** |  |  |  |  |  |
| Yes | 53 | 53 | 100 (93.24 – 100) | <0.001<br>(67.06) | ∞ |
| No | 482 | 197 | 40.87 (36.57 – 45.31) |  | 1 |
| **Tools for bat hunting** |  |  |  |  |  |
| Hands | 14 | 14 | 100 (78.24 – 100) | 0.07<br>(7.05) | ∞ |
| Stick | 99 | 67 | 67.67 (57.96 – 76.08) |  | 1.12 (0.43 – 2.90) |
| Others** | 14 | 11 | 78.57 (52.41 – 92.43) |  | 1.96 (0.42 – 9.11) |
| Chemicals | 23 | 15 | 65.21 (44.89 – 81.19) |  | 1 |
| **Localities** |  |  |  |  |  |
| Djalingo | 62 | 40 | 64.52 (52.09 – 75.26) | <0.001<br>(47.74) | 7.42 (3.28 – 16.82) |
| Lagdo | 70 | 44 | 62.86 (51.15 – 73.23) |  | 6.91 (3.12 – 15.32) |
| Babla | 87 | 51 | 58.62 (48.12 – 68.39) |  | 5.78 (2.70 – 12.39) |
| Garoua II | 70 | 35 | 50.00 (38.60 – 61.40) |  | 4.08 (1.86 – 8.96) |
| Mayo Oulo | 77 | 33 | 42.86 (32.41 – 53.99) |  | 3.06 (1.41 – 6.65) |
| Guider | 108 | 35 | 32.41 (24.32 – 41.71) |  | 1.96 (0.93 – 4.14) |
| Yelwa | 61 | 12 | 19.67 (11.63 – 31.31) |  | 1 |

\*\* includes the net, sling, clothes, branch of thorn. OR: Odds ratio; CI: confidence interval; \*: p
<0.05. N: number of answers to specific question

Additionally, respondents with bats in their homes (57.27%) also reported bats in different places including in the roofs of bedrooms (24.1%), trees (14.2%), abandoned buildings (2.2%) and warehouses (2.1%). The following tools were mainly used for bat hunting: sticks (66%), chemical products (15.33%), bare hands (9.33%) and other tools (9.33%) include nets, sling, thorns and clothes. The reasons for hunting the bats were due to their nuisance (69.04%), for consumption (12.5%), playing with bats (10.12%), sociocultural practice (2.97%), leisure (3.57%) and research (1.78%). Some of the respondents (17.60%) reported being bitten and scratched by bats when hunting the animals, however, none of them were vaccinated against rabies and none sought rabies post exposure prophylactic measures.

## Discussion

The present study reports the first serological evidence of Lyssavirus in bats in Cameroon and revealed rabies antigens in bats that appeared healthy. This observation underlines a potential risk for human communities living in contact with bats.

The prevalence of Lyssavirus reported in the present study is lower than the serological prevalence reported in other studies. A prevalence of 59.3% was reported in bat colonies in Spain [37], a prevalence of 50.34% of the virus in vampires bats (*Desmodus rotundus*) in Brazil [38]; and in Kenya a prevalence 40 to 67% was reported in colonies of *Eidolon helvum* and 29 to 46% in *Roussetus aegyptiacus* [17].

The findings of the present study are similar to those observed in moribund vampire bats (*Desmodus rotundus*) (27.6%) tested by IFA and PCR in northern Brazil [39]. Active infection has been reported in apparently healthy (0.1%) and moribund bats in Kenya (9%) [17, 40]. The prevalence of Lyssavirus in this study is also higher than values reported among bat species in the United States (9% to 10%) [40] Canada (7%) [36] Spain (3.3 to 10%) [38], Nigeria (19%) [19] and Kenya (9%) [17]. The difference in infection rates could be associated with the method of collection and the period of sample collection, which was conducted during the dry season. This period is characterized by food scarcity and elevated temperatures as well as a peak of rabies infection in bat populations in the study sites. The influence of the season on the occurrence of rabies in bats such as higher rates during the dry season [41] and fluctuating and cyclical rabies like infection in bat populations [40] were reported in other studies. The immigration of rabid bats into a colony can also cause a rapid spread of the infection through bites of infected bats during mutual grooming, mating and aggressive behaviour such as protection of the territory [42]. Unfavourable living conditions (climate, food source, immunity, density of the colony, parasite load among others), sex, age and physical depletion caused by stress, co-infections, sexual exhaustion and migration are also major risk factors for rabies infection [10, 17, 38, 40]. However, aggressive behaviour has been associated with a higher prevalence of rabies. Salmon-Mulanovich et al [43] reported a higher prevalence of rabies in male bats with more aggressive behaviour such as territory protection, defence against intrusion, fighting, and licking of body fluids during the breeding season. There was also a high or similar prevalence of Lyssavirus in female bats compared to males, which was associated with gregarious maternity behaviour during hibernation [44]. In the present study, young bats were less likely to be infected with rabies related virus than adults and older bats. This could be due to the fact that adult and older bats exhibited usual aggressiveness and adventurous behaviour for longer periods of time in the colonies interacting (continuous or intermittent) with other congeners in the same and other colonies (some of which may be infected) compared to young and juvenile bats.. In addition, serological prevalence can be influenced by antibodies of maternal origin and confer lasting immunity (6 to 8 weeks) to their offspring through placental and mammary routes [40], which could provide further protection to the young and juvenile bats.

Overall, the findings of this study agree with those of Salmon – Mulanovich *et al*. [43] in Peru, Kuzmin *et al*. [17] in Kenya and Dzikwi *et al*. [19] in Nigeria who reported that the location of the bats does not influence rabies infection rates in bats. This is in contrast to the findings of Costa *et al.* [45] who observed significant variation of bat Lyssavirus rates according to location. This variation could be associated to sampling biases such as disparities in distances between location sites of bat populations and the number of bats sampled at the different study sites [45]. Given the vast spatio-temporal spread of the activities of bat colonies, bat populations in close locations (less than 100km apart) will have common features since they frequently mix, interact together and could be considered as the same colony in this study.

The zoonotic transmission of bat Lyssavirus is well documented. Bat rabies can be transmitted to humans and other mammals over long periods and vast geographical areas through various routes including aerosols, unapparent bites, and contamination of nerve tissue with saliva, urine and other body fluids of infected bats [2, 3, 4, 5, 6, 7]. Based on the public health perception of bat rabies related virus, many respondents in rural localities in Garoua who shared the same environments with bat colonies were aware of canine and human rabies. This finding was significantly less than that observed by Bouli *et al.* [21] who reported over 88.73% awareness level of canine rabies in urban Garoua areas among respondents who were more literate and with better educational levels than in rural areas, as was the case in the present study. This is also in line with Moran *et al*. [35] who reported that over 91% of the respondents living in the rural areas harbouring bat populations in Guatemala had little or no knowledge of rabies. Costa and Fernandes [46] had also observed a strong positive correlation between educational levels and knowledge about rabies. Rural respondents with higher levels of education showed better knowledge of rabies in the present study.

Though the awareness about canine rabies was significantly relevant, most respondents seemed to have little knowledge of the potential zoonotic transmission of bat Lyssavirus. Irrespective of the respondents’ level of knowledge of canine rabies and bat Lyssavirus, the results showed that respondents who were aware of canine rabies and had bats in their homes and localities were more involved in bat hunting, and also interacted more with bats and bat products, compared to respondents who consume bats. Bouli *et al.*, [21] reported a lower awareness level of bat rabies (0.3%) in communities of Garoua-Cameroon, while higher levels ranging from 10% – 42% were reported in Thailand and Guatemala due to widespread public awareness campaigns against rabies in these countries [35, 47]. Similar to the level of knowledge about canine rabies found in the present study, the levels of awareness about bat Lyssavirus was significantly associated with literacy and educational levels and the age of the respondents. This finding highlights the lack of knowledge of the communities regarding the potential risk of bats Lyssavirus and its transmission to other species and humans. Awah-Ndukum et al., [23] reported that dogs and cats were the main source and transmitters of rabies in Cameroon, most likely due to the common and visible manifestations of the disease in these species and not in other species in the country.

The clinical manifestations of rabies are more furious and lethal in dogs [3] and non-lethal in bats [14]. The furious forms of rabies in dog usually constitute a canine rabies outbreak alert for human communities. Canine and human rabies is endemic in Cameroon, [21, 23]. In the present study, locality, age and educational level significantly influenced the levels of awareness of bat Lyssavirus and its zoonotic risk. Also, the presence of bats in homes, bat hunting and type of tools used for bat hunting as well as gender, religion and ethnicity of respondents played a major role on the level of human exposure to bats. Consumption of bats was frequent in the study sites, which is in agreement with Kamins *et al.* [48] who reported that bat consumption was cultural and a food habit in Ghana. This habit to hunt and consume bats as meat and source of animal protein among communities explains the difference in level of exposure observed in the present study between ethnic and religious groups. Indeed, according to the respondents in this study, Islam forbid eating some animals including bats while the other religions have not diet restriction.

Due to a technical problem impacting samples conservation (RNA degradation) the molecular part of this work could not be performed. It would be important to carry out an in-depth molecular study to accurately identify the type of Lyssavirus circulating in the region. This could help to understand the epidemiological pattern of Lyssavirus and identify potential reservoirs in Cameroon.

## Conclusion

Rabies is a zoonotic disease of all warm-blooded animals including man. Though the disease is endemic in Cameroon, with dogs being the main source and transmitters, bat habitats are widespread in the many Cameroon environments. Bats usually form colonies in the roofs, trees and abandoned tall structures in human communities and they are hunted as a source of animal protein in the the Northern region of Cameroon. The present study has shown a 26.89% prevalence of Lyssavirus.. This study is the first to report evidence of Lyssavirus in bats in Cameroon and revealed that rabies exists in apparently healthy bats and constitutes a human health problem in communities with bat habitats in the Northern region of the country. In order to isolate and determine the Lyssavirus genotype circulating in the bat population in Cameroon, future large-scale studies on bats should continue. Public awareness campaigns and health education are also essential to evaluate risk factors, develop protective measures against rabies and understand the potential role of bats as reservoirs of rabies especially in human communities where bat colonies are found. This study highlights the importance of investigating the dynamics of rabies virus transmission among animals and humans in this region and other parts of the country and to characterise the Lyssavirus type circulating among these flying mammals in Cameroon. Multi-sectorial sensitization of communities in Cameroon to improve their level of awareness on bat rabies and integration of the “One Health” approach for effective management of rabies in Cameroon should be emphasized.

## Acknowledgement

We are thankful to all the technicians in the virology laboratory section at Animal Pathology Department of LANAVET Garoua for facilitating our work in this study.

## Author’s contribution

**Conceptualization**: Rodrigue Simonet Poueme Namegni, Isaac Dah, Julius Awah-Ndukum

**Data curation**: Moctar Mouiche Mouliom Mohamed, Julius Awah-Ndukum Ranyl Nguena Guefack Noumedem, Isaac Dah,

**Formal analysis**: Julius Awah-Ndukum, Ranyl Nguena Guefack Noumedem, Isaac Dah,

**Investigation:** Isaac Dah, Laurent God-Yang, Rodrigue Simonet Poueme Namegni

**Methodology**: Isaac Dah, Rodrigue Simonet Poueme Namegni, Moctar Mouiche Mouliom Mohamed, Simon Dickmu Jumbo, Isabelle Conclois.

**Resources:** Rodrigue Simonet Poueme Namegni, Simon Dickmu Jumbo, Abel Wade.

**Software:** Isaac Dah, Ranyl Nguena Guefack Noumedem, Laurent God-Yang, Jean Marc Feussom Kameni

**Supervision**: Dorothee Missé, Liegeois Florian, Rodrigue Simonet Poueme Namegni, Abel Wade, Jean Marc Feussom Kameni, Julius Awah-Ndukum

**Validation**: Liegeois Florian, Dorothee Misse, Abel Wade, Julius Awah-Ndukum

**Writing original draft preparation**: Isaac Dah, Rodrigue Simonet Poueme Namegni Julius Awah-Ndukum

**Writing – review & editing**: Isaac Dah, Rodrigue Simonet Poueme Namegni, Moctar Mouiche Mouliom Mohamed, Liegeois Florian, Abel Wade, Dorothee Missé, Julius Awah-Ndukum

All the authors have approved the final version of manuscript and agreed for submission

## Availability of data

The datasets used and/or analysed during the current study are available from the corresponding author on reasonable request

